# Development and Validation of a Nomogram Prognostic Model for Small-Cell Lung Cancer Patients

**DOI:** 10.1101/273029

**Authors:** Shidan Wang, Lin Yang, Matthew Maclean, David E. Gerber, Guanghua Xiao, Yang Xie

## Abstract

**Background:** Small-cell lung cancer (SCLC) accounts for almost 15% of lung cancer cases in the United States. Nomogram prognostic models could greatly facilitate risk stratification and treatment planning, as well as more refined enrollment criteria for clinical trials. We developed and validated a new nomogram prognostic model for SCLC patients using a large SCLC patient cohort from the National Cancer Database (NCDB).

**Methods:** Clinical data of 24,680 SCLC patients diagnosed from 2004 to 2011 were used to develop the nomogram prognostic model. The model was then validated using an independent cohort of 9,700 SCLC patients diagnosed from 2012 to 2013. The prognostic performance was evaluated using p value, concordance index and integrated Area Under the (time dependent Receiver Operating Characteristic) Curve.

**Results:** The following variables were contained in the final prognostic model: age, gender, race, ethnicity, Charlson/Deyo Score, TNM Stage (assigned according to the AJCC 8^th^ edition), treatment type (combination of surgery, radiation therapy and chemotherapy), and laterality. The model was validated in an independent testing group with a concordance index of 0.722 ± 0.004 and an integrated AUC of 0.79. The nomogram model has a significantly higher prognostic accuracy than previously developed models, including the AJCC 8^th^ edition TNM-staging system. We implemented the proposed nomogram and two previously published nomograms in an online webserver.

**Conclusions:** We developed a nomogram prognostic model for SCLC patients, and validated the model using an independent patient cohort. The nomogram performs better than earlier models, including AJCC staging.

## Introduction

Lung cancer is the leading cause of death from cancer in the United States and worldwide. Small-cell lung cancer (SCLC) accounts for 13.6% of all lung cancer cases (1, 2). Compared to non-small-cell lung cancer (NSCLC), in which the 5-year survival rate is 18.0%, SCLC has only a 6.2% 5-year survival rate, and is characterized by a more rapid tumor growth rate and death from recurrent disease (3, 4). Over the last several decades, there have been only modest improvements in patient survival (5) and no molecularly targeted therapy has proven beneficial for SCLC patients (3). Nomogram prognostic models that predict patient outcomes may facilitate better treatment stratification and outcome evaluation, as well as more refined patient enrollment criteria for clinical trials in SCLC. Furthermore, a recent study in breast cancer (6) showed that user-friendly online prognostic tools could greatly enhance patient care. However, currently there are no such online tools available for prognosis of SCLC.

To date there is only one study of nomograms in SCLC, published by Xie et al (4), which developed two nomograms: one for extensive stage SCLC patients and the other for limited stage SCLC patients. The nomograms developed from that study provide useful tools for clinicians and researchers to stratify the risk of SCLC patients. However, the study simply classified patients as limited or extensive stage without using the more accurate TNM staging proposed by the International Association for the Study of Lung Cancer (IASLC) (7). Furthermore, there is a lack of independent validation for this model, probably due to the limited sample size (n = 555 for extensive stage and n = 383 for limited stage). Other non-nomogram prognostic models include the Manchester score and Spain score. However, both of these were developed on small sample sets (n = 407 for Manchester score and n = 341 for Spain score) and divide patients into only three risk groups (8, 9).

The goal of this study was to identify prognostic factors for SCLC patients, and then develop and validate a new nomogram prognostic model in a large SCLC patient cohort. The National Cancer Database (NCDB) includes over 200,000 patients diagnosed with SCLC from 2004 to 2013 in the United States, of which 34,380 SCLC patients without any missing values were used to develop and validate our nomogram prognostic model. The SCLC cases in the NCDB dataset were separated into a training cohort and a validation cohort based on the year of diagnosis. The model was developed from the training cohort of 24,680 SCLC patients diagnosed from 2004 to 2011, and then validated in the validation cohort of 9,700 SCLC patients diagnosed from 2012 to 2013. The prognostic performance was evaluated using p value, concordance index and integrated Area Under the Curve. In order to facilitate public usage, we implemented our nomogram and the previous ones by Xie et al. in an online webserver. Compared to the previously published models, our model has the following advantages: 1) it was validated in an independent set; 2) it was developed and validated with a much larger sample size; 3) it was developed across multiple facilities and facility types, which greatly diminishes sample selection bias; 4) it utilizes accurate SCLC staging criteria: the AJCC 8^th^ edition TNM staging system proposed by IASLC (10); and 5) it provides an online webserver so that clinicians can use the nomogram model easily.

## Methods

### Source of data

202,194 SCLC cases were identified from NCDB and 34,380 of them met our inclusion criterion that they do not contain any missing data for selected variables. The source of missing values is listed in **Supplementary Table 1**. The cases are independent and recorded by annual reports from all the CoC-accredited programs from 2004 to 2013. 24,680 cases that were diagnosed from 2004 to 2011 were assigned to the training group and used to develop the nomogram prognostic model. The 9,700 cases diagnosed from 2012 to 2013 were assigned to the testing group and used to validate the model.

### Nomogram development

The nomogram was developed using the training cohort of 24,680 patients diagnosed from 2004 to 2011. Overall survival was used as the primary outcome. Two extra variables were first constructed based on NCDB variables: treatment was defined as the stratification result of surgery, chemotherapy and radiation therapy; TNM stage was defined according to the coding guidelines of the Collaborative Staging Manual and Coding Instructions for the new 8^th^ edition lung cancer staging system defined by the American Joint Committee on Cancer (AJCC) and the Union for International Cancer Control (UICC) (11–14), and followed Yang et al’s method (15). The input variables were age, gender, race, Hispanic origin, Charlson/Deyo Score, sequence number, primary site, laterality, grade, 8^th^ edition TNM stage and treatment type.

Univariate Cox regression and Wald test were then used to screen for variables that were significantly correlated with overall survival in the training group. Predictors with a p-value less than 0.05 were fed to a multivariate Cox regression model. Backward stepwise selection based on Akaike Information Criteria (AIC) was used to further eliminate redundant variables. The resulting multivariate Cox regression model was used to calculate risk score and build the final nomogram prognostic model.

### Model validation

To validate our model, three criteria were used to evaluate prediction performance in the testing set. First, the cases were grouped according to their predicted risk score, and Kaplan-Meier survival curves and Wald test were used to compare survival differences among the groups. Second, a concordance index (c-index) was calculated to estimate the similarity between the ranking of true survival time and of predicted risk score. Third, area under the curve (AUC) of time-dependent receiver operating characteristics (ROC) (16, 17) was calculated at each month from the 1^st^ to the 30^th^ month. Integrated AUC was calculated by averaging the 30 AUC values.

The other two models, the AJCC 8^th^ edition TNM staging system and the traditional limited/extensive staging system, were also tested for prognostic performance in the testing group. C-index and integrated AUC were used to compare this nomogram with the two staging systems. Here, extensive stage was defined based on the presence of distant metastases (M1 stage) (18, 19). All other cases (M0 stage) were grouped as limited stage.

All computations were conducted in the R environment, version 3.3.2 (20). R packages “survival” (version 2.40-1) and “timeROC” (version 0.3) were used. Results with p-value ≤ 0.05 were considered statistically significant.

### Implementation of this and previously published models

To facilitate researchers’ and clinicians’ usage of our model, we created a user-friendly webserver for our nomogram and the two models from Xie et al (4). The nomogram from this study calculates the risk score, plots the survival curve and provides survival probabilities for 120 months at 6-month increments. The Xie et al models for both extensive and limited stage cases provide 6-month and 12-month survival probabilities and predicted median survival time. Data points were read from Figures 1 and 2 of the Xie et al publication (4), and the corresponding survival probability for a given score was calculated by linear interpolation.

**Figure 1.**
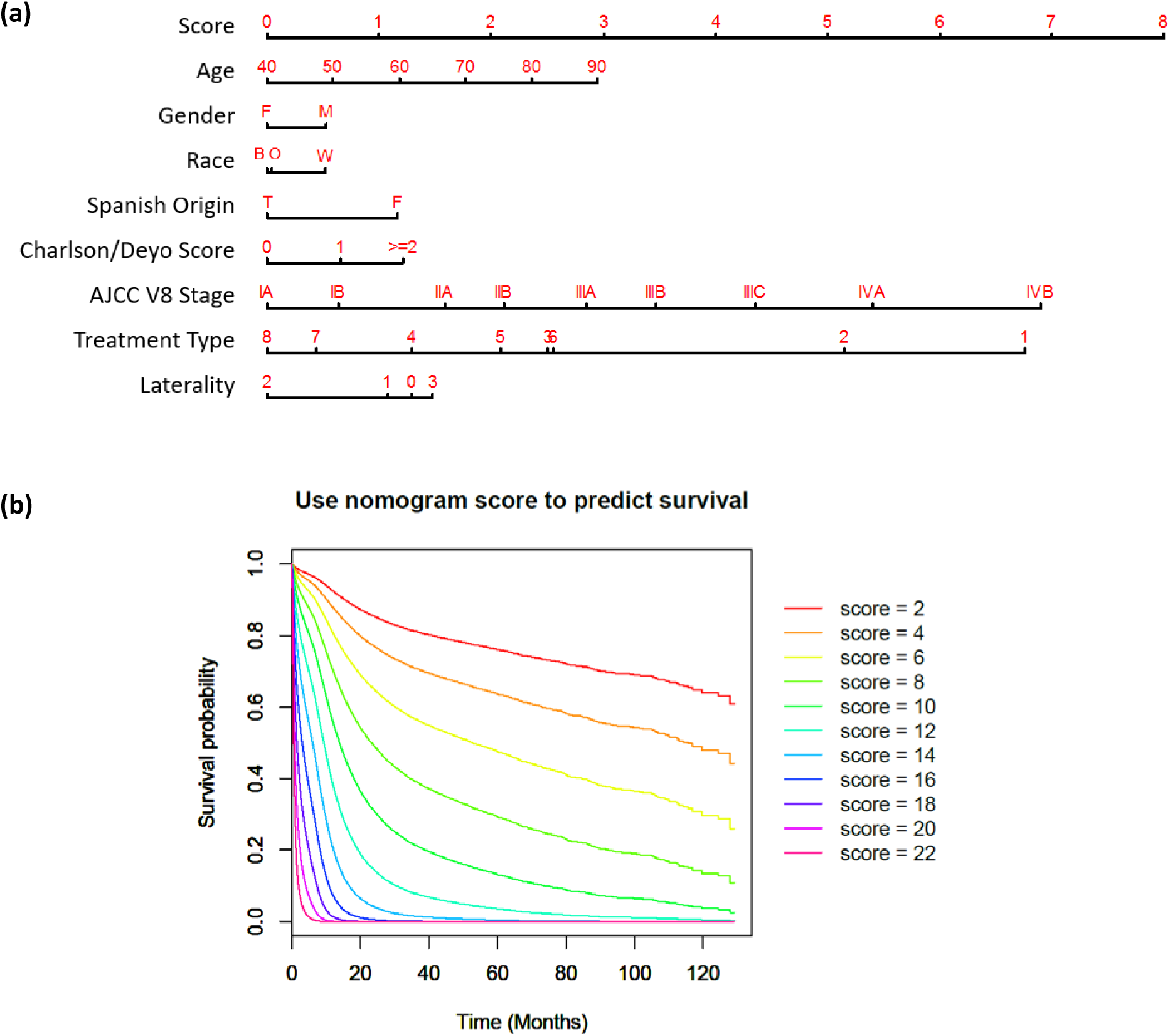
Nomogram to calculate risk score and predict survival probability. **(a)** Race includes black (B), white (W) and other (O). Treatment types include: no surgery, no chemo, no radiation (1); no surgery, no chemo, radiation done (2); no surgery, chemo done, no radiation (3); no surgery, chemo done, radiation done (4); surgery done, no chemo, no radiation (5); surgery done, no chemo, radiation done (6); surgery done, chemo done, no radiation (7); and surgery done, chemo done, radiation done (8). Laterality of tumor origin includes: not a paired site (0), only one side (either left or right) is involved (1), bilateral involvement (2), paired site with unknown origin side or midline tumor (3). **(b)** Predicted patient survival probability curve corresponding to risk scores ranging from 2 to 22.

**Figure 2.**
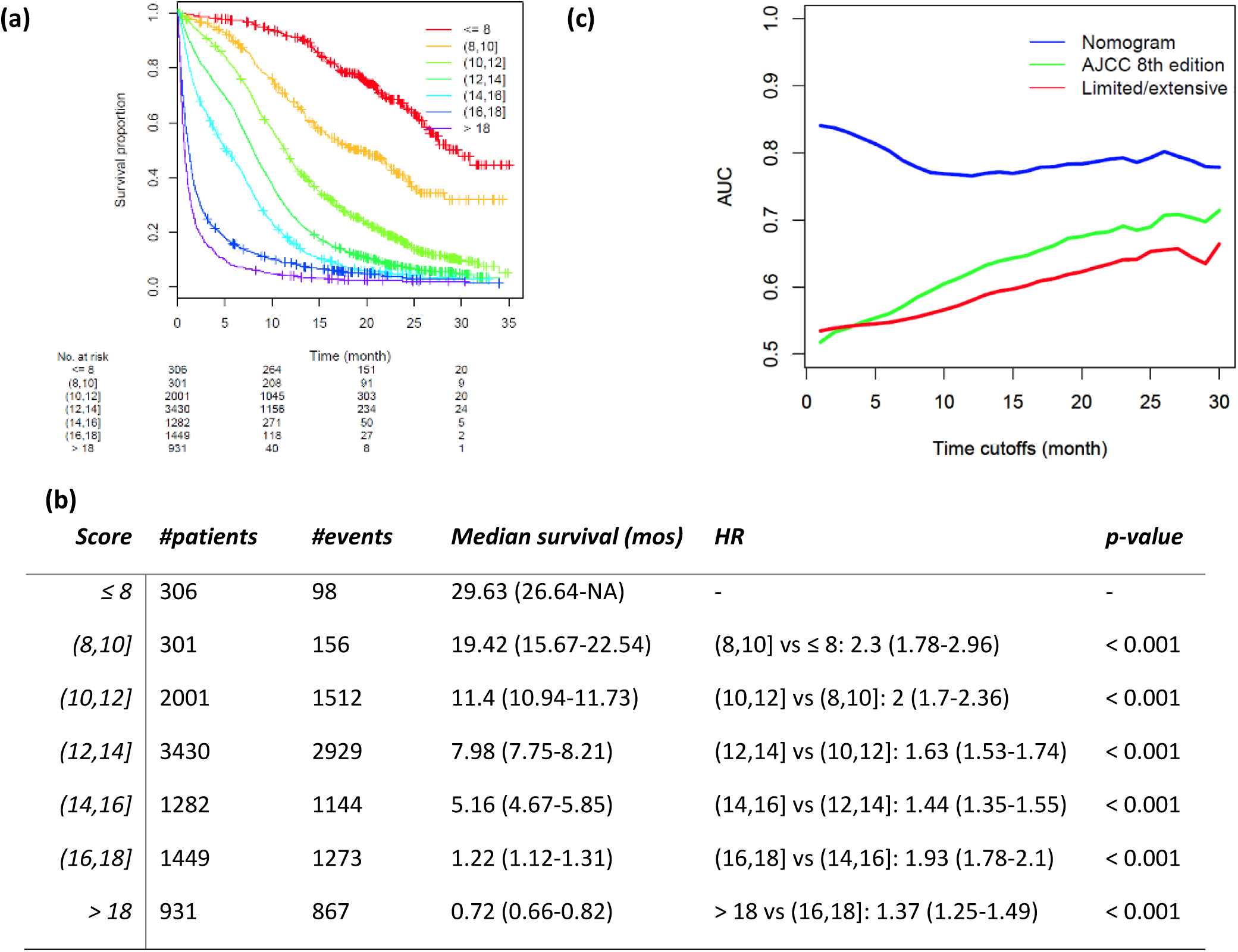
Validation of proposed nomogram prognostic model in the testing set. **(a)** Risk scores of testing set cases were calculated according to the model in Figure 1 and grouped into 8 subgroups. K-M plot was depicted for each group. **(b)** Summary of groups in (a). Hazard Ratio (HR) was calculated using Coxph regression model between each two adjacent lines. P-value was calculated using Wald test. **(c)** Area under the curve (AUC) was calculated for three prognostic models for every month from the 1^st^ to the 30^th^ month. Blue: nomogram developed in this study; green: AJCC 8^th^ TNM staging system; red: limited/extensive staging system.

## Results

### Characteristics of the training and validation cohorts

In total, 202,194 SCLC cases were identified in NCDB, among which, 34,380 cases that did not contain any missing variables were included in this study. Based on year of diagnosis, included cases were divided into two distinct groups: cases that were diagnosed from 2004 to 2011 (n = 24,680) were used as the training cohort, while cases that were diagnosed from 2012 to 2013 (n = 9,700) were used as the validation cohort. Characteristics of the two sets are shown in **Table 1**. Demographic variables were similar between training and testing sets.

**Table 1.** Characteristics of training set and testing set. P-values were calculated by Chi-square test.

|  | *training set (%)* | *testing set (%)* | *p-value* |
| --- | --- | --- | --- |
| *No. of cases* | 24,680 | 9,700 |  |
| *Year of diagnosis* | 2004-2011 | 2012-2013 |  |
| *Age* |  |  | 0.09 |
| < 65y | 9,559 (38.7) | 3,855 (39.7) |  |
| ≥ 65y | 15,121 (61.3) | 5,845 (60.3) |  |
| *Gender* |  |  | 0.9 |
| Male | 12,240 (49.6) | 4,803 (49.5) |  |
| Female | 12,440 (50.4) | 4,897 (50.5) |  |
| *Race* |  |  | 0.73 |
| White | 22,276 (90.3) | 8,779 (90.5) |  |
| Black | 1,912 (7.7) | 727 (7.5) |  |
| Other | 492 (2) | 194 (2) |  |
| *Ethnicity* |  |  | 0.91 |
| Non-Hispanic | 24,084 (97.6) | 9,463 (97.6) |  |
| Hispanic | 596 (2.4) | 237 (2.4) |  |
| *Charlson/Deyo score* |  |  | <0.001 |
| 0 | 13,288 (53.8) | 5,031 (51.9) |  |
| 1 | 7,629 (30.9) | 3,061 (31.6) |  |
| ≥ 2 | 3,763 (15.2) | 1,608 (16.6) |  |
| *AJCC V8 TNM stage* |  |  | <0.001 |
| IA | 1,207 (4.9) | 160 (1.6) |  |
| IB | 463 (1.9) | 74 (0.8) |  |
| IIA | 140 (0.6) | 18 (0.2) |  |
| IIB | 853 (3.5) | 97 (1) |  |
| IIIA | 1,548 (6.3) | 156 (1.6) |  |
| IIIB | 902 (3.7) | 89 (0.9) |  |
| IIIC | 208 (0.8) | 27 (0.3) |  |
| IVA | 14,699 (59.6) | 6,655 (68.6) |  |
| IVB | 4,660 (18.9) | 2,424 (25) |  |
| *Surgery* |  |  | <0.001 |
| No | 21,824 (88.4) | 9,256 (95.4) |  |
| Yes | 2,856 (11.6) | 444 (4.6) |  |
| *Chemotherapy* |  |  | 0.49 |
| No | 7,175 (29.1) | 2,857 (29.5) |  |
| Yes | 17,505 (70.9) | 6,843 (70.5) |  |
| *Radiation therapy* |  |  | <0.001 |
| No | 14,549 (59) | 5,967 (61.5) |  |
| Yes | 10,131 (41) | 3,733 (38.5) |  |
| *Laterality* |  |  | <0.001 |
| Not a paired site | 2,298 (9.3) | 911 (9.4) |  |
| Only one side involved | 20,447 (82.8) | 8,016 (82.6) |  |
| Bilateral involvement | 624 (2.5) | 154 (1.6) |  |
| Paired site but lateral origin unknown;<br>midline tumor | 1,311 (5.3) | 619 (6.4) |  |

### Building nomogram prognostic model in training cohort

In univariate analysis, age, gender, race, Hispanic origin, Charlson/Deyo score, TNM stage by AJCC 8^th^ edition, treatment type, primary site, laterality, and grade were significantly associated with overall survival in the training group **(Table 2)**. After stepwise selection to further remove potential redundancy, age, sex, race, ethnicity, Charlson/Deyo score, TNM stage by AJCC 8^th^ edition, treatment type, and laterality were used in the final nomogram model (coefficients summarized in **Table 3**). The final risk score was calculated by adding up the score of each item using the nomogram depicted in **Figure 1a**. The TNM stage defined by the AJCC 8^th^ edition contributed the most to the survival prognosis, followed by the treatment type and age. The predicted survival probability using the Cox regression model of risk scores was plotted in **Figure 1b**.

**Table 2.** Univariate analysis results summary. HR: Hazard Ratio, CI: Confidence Interval.

| *Variable* | *HR (95% CI)* | *p-value* |
| --- | --- | --- |
| *Age* | 1.023 (1.023-1.024) | < 0.001 |
| *Sex (Female vs. Male)* | 0.84 (0.83-0.85) | < 0.001 |
| *Race* |  |  |
| *White* | 1 (reference) | - |
| *Black* | 0.97 (0.95-0.99) | 0.006 |
| *Other* | 0.94 (0.90-0.97) | 0.001 |
| *Hispanic origin (Yes vs. No)* | 0.95 (0.92-0.99) | 0.028 |
| *Charlson/Deyo score* |  |  |
| 0 | 1 (reference) | - |
| 1 | 1.22 (1.20-1.24) | < 0.001 |
| ≥ 2 | 1.59 (1.56-0.61) | < 0.001 |
| *Sequence Number* |  |  |
| 0 | 1 (reference) | - |
| 1 | 1 (0.98-1.01) | 0.82 |
| ≥ 2 | 1.01 (0.92-1.11) | 0.83 |
| *AJCC V8 TNM Stage* |  |  |
| IA | 1 (reference) | - |
| IB | 1.22 (1.07-1.39) | < 0.001 |
| IIA | 1.63 (1.34-1.98) | < 0.001 |
| IIB | 1.6 (1.45-1.78) | < 0.001 |
| IIIA | 2.12 (1.94-2.31) | < 0.001 |
| IIIB | 2.55 (2.32-2.81) | < 0.001 |
| IIIC | 3.26 (2.81-3.78) | < 0.001 |
| IVA | 5.25 (4.88-5.65) | < 0.001 |
| IVB | 7.04 (6.51-7.61) | < 0.001 |
| *Surgery (With vs. Without)* | 0.39 (0.38-0.4) | < 0.001 |
| *Chemotherapy (With vs. Without)* | 0.38 (0.37-0.38) | < 0.001 |
| *Radiation therapy (With vs. Without)* | 0.52 (0.51-0.52) | < 0.001 |
| *Primary Site* |  |  |
| C340 | 1 (reference) | - |
| C341 | 0.89 (0.88-0.91) | < 0.001 |
| C342 | 0.9 (0.87-0.92) | < 0.001 |
| C343 | 0.96 (0.94-0.98) | < 0.001 |
| C348 | 1.07 (1.03-1.11) | < 0.001 |
| C349 | 1.13 (1.11-1.16) | < 0.001 |
| *Laterality* |  |  |
| *Not a paired site* | 1 (reference) | - |
| *Only one side involved* | 0.94 (0.93-0.96) | < 0.001 |
| *Bilateral involvement* | 1.47 (1.4-1.54) | < 0.001 |
| *Paired site but lateral origin unknown; midline tumor* | 1.18 (1.15-1.21) | < 0.001 |
| *Grade* |  |  |
| 1 | 1 (reference) | - |
| 2 | 0.99 (0.86-1.14) | 0.87 |
| 3 | 1.29 (1.15-1.46) | < 0.001 |
| 4 | 1.39 (1.23-1.56) | < 0.001 |

**Table 3.** Hazard Ratio (HR) and 95% confidence interval of nomogram parameters.

|  | HR (95% CI) | p-value |
| --- | --- | --- |
| Age | 1.01 (1.01-1.02) | < 0.001 |
| Sex (Female vs. Male) | 0.88 (0.85-0.9) | < 0.001 |
| Race |  |  |
| White | 1 (reference) | - |
| Black | 0.88 (0.84-0.92) | < 0.001 |
| Other | 0.89 (0.8-0.98) | 0.02 |
| Hispanic origin (Yes vs. No) | 0.75 (0.68-0.82) | < 0.001 |
| Charlson/Deyo score |  |  |
| 0 | 1 (reference) | - |
| 1 | 1.18 (1.14-1.21) | < 0.001 |
| >= 2 | 1.36 (1.31-1.41) | < 0.001 |
| AJCC V8 TNM stage |  |  |
| IA | 1 (reference) | - |
| IB | 1.17 (1.02-1.35) | 0.02 |
| IIA | 1.49 (1.2-1.84) | < 0.001 |
| IIB | 1.7 (1.52-1.9) | < 0.001 |
| IIIA | 2.04 (1.83-2.26) | < 0.001 |
| IIIB | 2.38 (2.11-2.68) | < 0.001 |
| IIIC | 2.97 (2.5-3.54) | < 0.001 |
| IVA | 3.86 (3.48-4.27) | < 0.001 |
| IVB | 5.62 (5.06-6.24) | < 0.001 |
| Treatment |  |  |
| No surgery, no chemo, no radiation | 1 (reference) | - |
| No surgery, no chemo, radiation done | 0.67 (0.63-0.71) | < 0.001 |
| No surgery, chemo done, no radiation | 0.35 (0.33-0.36) | < 0.001 |
| No surgery, chemo done, radiation done | 0.25 (0.24-0.26) | < 0.001 |
| Surgery done, no chemo, no radiation | 0.31 (0.28-0.35) | < 0.001 |
| Surgery done, no chemo, radiation done | 0.35 (0.27-0.46) | < 0.001 |
| Surgery done, chemo done, no radiation | 0.21 (0.19-0.23) | < 0.001 |
| Surgery done, chemo done, radiation done | 0.18 (0.17-0.2) | < 0.001 |
| Laterality |  |  |
| Not a paired site | 1 (reference) | - |
| Only one side involved | 0.95 (0.91-0.99) | 0.02 |
| Bilateral involvement | 0.72 (0.66-0.79) | < 0.001 |
| Paired site but lateral origin unknown; midline tumor | 1.05 (0.98-1.13) | 0.19 |

### Validation in testing cohort

The proposed nomogram was validated in the independent testing set (n=9,700). The survival difference between any two adjacent groups, which were grouped by predicted risk score, was significant (p-value < 0.05, **Figure 2a & ab**). The median survival times of score groups ranged from 0.7 months (when risk score > 18) to 30.9 months (when risk score < 6). The c-index was 0.722 ± 0.004 and the integrated AUC was 0.79 from the 1^st^ month to the 30^th^ month (**Figure 2c, Supplementary Table 2**).

With regards to prognostic ability, the proposed nomogram performed better than the two commonly used SCLC staging systems, the AJCC TNM system and limited/extensive staging system (**Figure 2c, Supplementary Table 2, Supplementary Figure 1 a&b**). The AUC of the nomogram was the highest throughout the 1^st^ to the 30^th^ month, followed by the 8^th^ edition TNM staging system. The integrated AUC of the proposed nomogram was 0.789, while those of the 8^th^ edition TNM staging system and the limited/extensive staging system were 0.634 and 0.598, respectively. The c-index of this nomogram (0.722 ± 0.004) was also significantly higher than the c-indexes of the 8^th^ edition TNM staging system (0.550 ± 0.003) and the limited/extensive staging system (0.539 ± 0.002), confirming the strong prognostic power of this proposed nomogram.

### Development of webserver for easy access of our own and previously published models

An online version of our nomogram (**Figure 3a**) can be accessed at http://lce.biohpc.swmed.edu/lungcancer/sclc_nomogram, to assist researchers and clinicians. Online implementation of the two nomograms from Xie et al (4) are also available (**Figure 3b&c**). Predicted survival probability across time can be easily determined by inputting clinical features and reading output figures and tables generated by the webserver.

**Figure 3.**
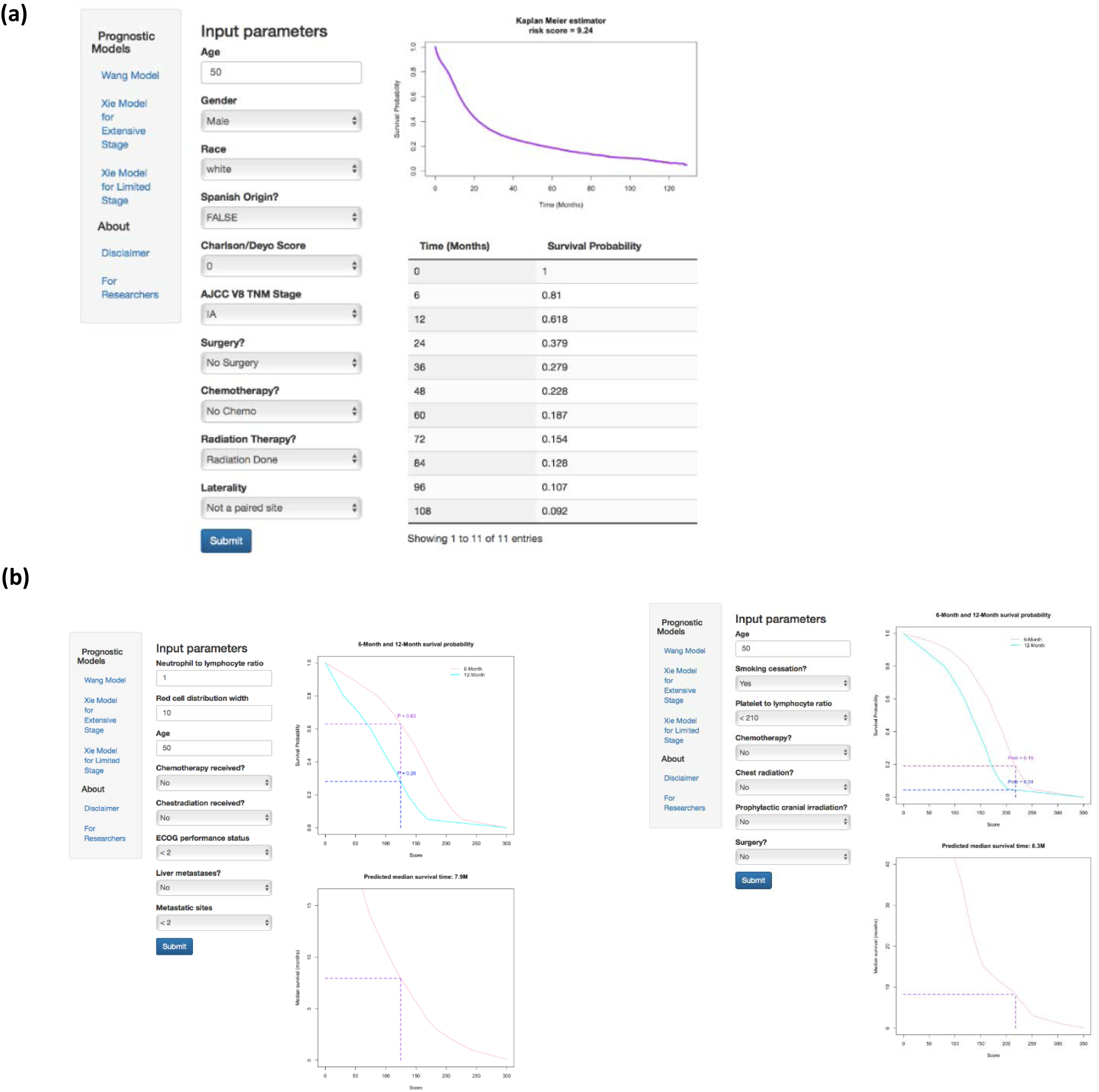
Online webserver interface for our nomogram as well as previous prognostic models. (a) The newly developed nomogram in this study (Wang Model). (b) 2 published models by Xie et al (Xie Model for Extensive Stage and Xie Model for Limited Stage)

## Discussion

In this study, a nomogram prognostic model was developed and validated using a large cohort of SCLC cases across the United States. This nomogram, based on routinely available demographic, staging and treatment information, predicts the survival probability for individual SCLC patients. The publicly accessible online implementation will assist clinicians in making treatment decisions.

Compared with other prognostic indexes, such as the Manchester Score (8) and the Spain prognostic index (9), our model calculates individualized survival probability rather than assigning cases into a few risk groups, thus better capturing heterogeneity across patients. Compared with the previously published nomogram by Xie et al., this model used a much larger training dataset and involved multiple treating facilities, which allowed for smaller sampling bias. The internal c-index of this model was 0.744 ± 0.002, higher than the previously published models (0.73 for both nomograms in (4)). Independent validation of our model showed significantly different outcomes among different score groups (**Figure 2a&b**). High concordance index (0.722 ± 0.004) and integrated AUC score (0.789, **Figure 2c, Supplementary Table 2**) in the testing set also indicated the strong predictive ability of our nomogram model. In addition, combining demographic, clinical and treatment information together produced a nomogram with better performance than using staging information alone (**Figure 2b, Supplementary Table 2**). Thus, this comprehensive and individualized risk score calculation method could be used as stratification criteria in randomized studies and clinical trials.

In this nomogram, age, gender, race, ethnicity, Charlson/Deyo score, AJCC 8^th^ edition stage, treatment type and laterality were kept after univariate Cox regression screening and backward stepwise selection. Age, gender, and Charlson/Deyo score have previously been shown significantly relevant to survival of SCLC patients (4, 21). Noticeably, AJCC 8^th^ edition stage contributed the most to the final risk score (**Figure 1a**), with clear distinctions between each two adjacent TNM stages (**Table 3**), and showed better prognostic performance than limited/extensive staging system with higher c-index and AUC (**Figure 2b, Supplementary Table 2**). The significant contribution of TNM stage to this nomogram externally validates the performance of the 8^th^ edition TNM lung cancer classification system, and highlights the importance of applying this more accurate staging system to SCLC rather than using the traditional limited/extended staging (7, 10, 22).

This proposed nomogram also illustrates the prognostic implications of using different treatment methods (**Figure 1a, Table 3**). As expected, cases treated with both surgery and chemo-radiation therapy have the lowest risk score and cases not treated with any method have the highest risk score. Furthermore, the nomogram (Figure 1b) is consistent with current research in that it predicts better survival for surgery with chemo-radiation (treatment type 7 and 8) than for surgery with chemotherapy alone (type 3 and 4) [21]. However, the risk scores of different treatment methods are not recommended to be directly used as a guideline for treatment selection, since clinical treatment decisions should be made based on multiple factors such as TNM stage and patient comorbidities(3).

There were several limitations in the development of this nomogram. The first limitation was a lack of some routinely available clinical data, such as the neutrophil to lymphocyte ratio (NLR) and platelet to lymphocyte ratio (PLR). The absence of this information prevented direct comparison of performance between our model and another published nomogram (4). Constructing a prognostic model using both the factors identified in our model and other lab tests such as NLR would thus be beneficial in creating an even more accurate prognostic prediction. The second limitation was the inability to capture interaction terms among the predictors. For example, patients with early stage disease (stage I & II) were more likely to receive surgery than patients with late stage disease (stage III and IV). To satisfy the requirement for convenience and interpretability of the nomogram, interaction terms were not considered in this model. However, a more complex model considering all potential interaction terms would be expected to have better prognostic performance. Finally, out of 200,000 SCLC patients from the NCDB, there are only 34,380 patients without missing values. This large percent of missing data might introduce some selection bias.

## Conclusion

We developed a nomogram prognostic model for SCLC patients, and validated the model using an independent patient cohort. The proposed nomogram shows better prognostic performance than other existing models. This nomogram and previously published prognostic models were implemented on an online webserver. Researchers, clinicians and patients can easily predict the survival probability for each individual patient using this webserver.

## Acknowledgements

This work was supported by the National Institutes of Health [5R01CA152301, P50CA70907, 5P30CA142543, 1R01GM115473, K24CA201543 and 1R01CA172211]; the Cancer Prevention and Research Institute of Texas [RP120732]

The NCDB is a joint project of the Commission on Cancer of the American College of Surgeons and the American Cancer Society. The data used in this study is derived from a de-identified NCDB file. The American College of Surgeons and the Commission on Cancer have not verified and are not responsible for the analytic or statistical methodology employed, or the conclusions drawn from this data by the investigator.

## Supplementary Material

**Supplementary Table 1.** Source of missing data.

| *Missing characteristics* | *Number of cases* | *% of total excluded cases* |
| --- | --- | --- |
| *Any* | 167,814 | 100 |
| *Hispanic origin* | 15,813 | 9.4 |
| *8<sup>th</sup> AJCC TNM stage* | 154,465 | 92.0 |
| *8<sup>th</sup> AJCC T stage* | 70,101 | 41.8 |
| *8<sup>th</sup> AJCC N stage* | 157,297 | 93.7 |
| *8<sup>th</sup> AJCC M stage* | 86,441 | 51.5 |
| *Treatment method* | 4,166 | 2.5 |

**Supplementary Table 2.**
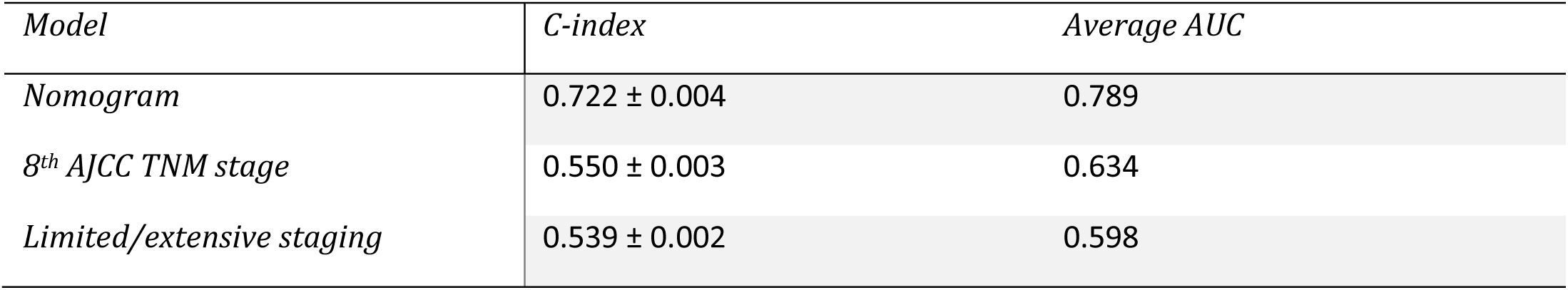
Comparison of prognostic performance of three models in testing dataset.

**Supplementary Figure 1.**
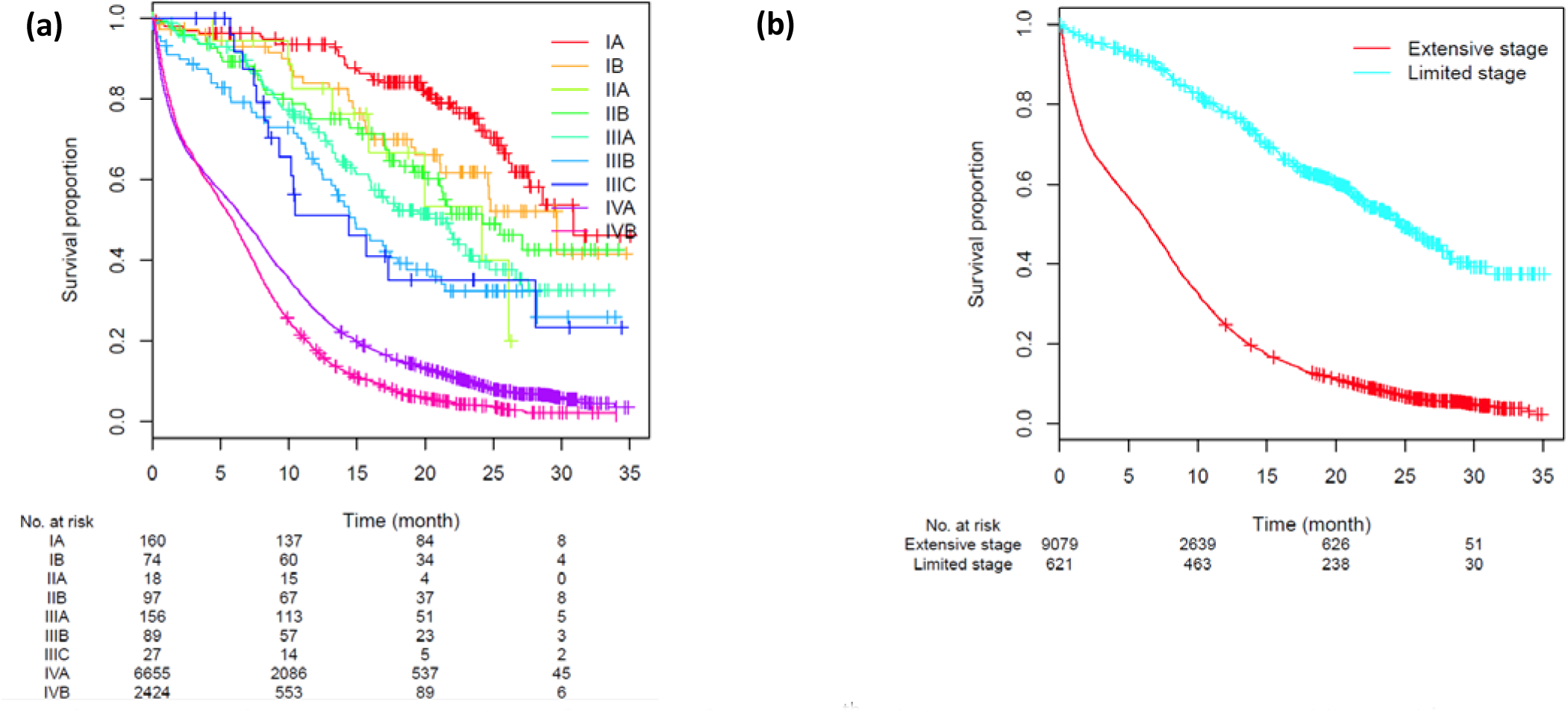
Prognostic performance for AJCC 8^th^ edition TNM staging system and limited/extensive staging system. (a) K-M plot grouped by AJCC 8^th^ edition TNM stages. (b) K-M plot grouped by limited/extensive stages.

